# Inulin prebiotic reinforces host cancer immunosurveillance via γδ T cell activation

**DOI:** 10.1101/2022.10.13.512019

**Authors:** Emilie Boucher, Caroline Plazy, Mathias L. Richard, Antonia Suau, Irène Mangin, Muriel Cornet, Delphine Aldebert, Bertrand Toussaint, Dalil Hannani

## Abstract

The gut microbiota is now recognized as a key parameter affecting the host’s anti-cancer immunosurveillance and ability to respond to immunotherapy. Therefore, optimal modulation for preventive and therapeutic purposes is very appealing. Diet is one of the most potent modulators of microbiota, and thus nutritional intervention could be exploited to improve host anti-cancer immunity. Here, we show that an inulin-enriched diet, a prebiotic known to promote immunostimulatory bacteria, triggers an enhanced Th1-polarized CD4^+^ and CD8^+^ αβ T cell-mediated anti-tumor response and attenuates tumor growth in three preclinical tumor-bearing mouse models. We highlighted that the inulin-mediated anti-tumor effect relies on the activation of both intestinal and tumor-infiltrating γδ T cells that are indispensable for αβ T cell activation and subsequent tumor growth control, in a microbiota-dependent manner. Overall, our data identified these cells as a critical immune subset, mandatory for inulin-mediated anti-tumor immunity *in vivo*, further supporting and rationalizing the use of such prebiotic approaches, as well as the development of immunotherapies targeting γδ T cells in cancer prevention and immunotherapy.

**Significance:** Our study reveals that γδ T cells anti-cancer activity can be improved by nutritional intervention, in a microbiota-dependent manner. This work also indicates that γδ T cells are indispensable for reinforcing αβ T cells cancer immunosurveillance and subsequent tumor growth control. We believe that these findings could be of interest to the field of gut microbiota modulation, rationalizing the use of such prebiotic approaches as well as γδ T cells targeting, in cancer prevention and immunotherapy.

## Introduction

According to the World Health Organization (WHO), cancer is the second leading cause of death globally, and the increasing cancer incidence is a major public health issue worldwide (1). Despite the development of innovative treatments, such as immunotherapy, including immune checkpoint blockers (ICB), which have recently revolutionized patient care, quality of life, and survival, huge efforts are still needed to prevent and treat cancers. It is worth noting that between 30% and 50% of cancer deaths can be prevented by modifying or avoiding major risk factors and implementing existing evidence-based prevention strategies, including eating a healthy diet, primarily fruits and vegetables (2). A healthy diet helps prevent the development of cancer in several ways that may be synergistic. It limits host cell oxidation and metabolism as well as genetic instability and chronic inflammation. It also affects gut microbiota composition and functions, and subsequently, host immune functions (3).

Indeed, the host immune tonus is a key parameter in cancer prevention through immunosurveillance, which relies on the ability of different immune cell subsets to patrol, recognize, and eliminate transformed cells (4). Both innate and adaptive immunity participate in this process and prevent cancer development by recognizing stress-induced ligands, dysregulation of tumor cell metabolism, or peptides derived from tumor antigens. Effector CD8^+^ T cells are among the most important subsets involved in cancer surveillance and are key targets in immunotherapy. Furthermore, γδ T cells are unconventional T cells representing a unique population displaying multiple functions, including potent anti-tumor activity (5,6). Indeed, γδ T cells are key players in cancer immunosurveillance, as they can recognize transformed cells expressing stress-ligands via innate receptors (e.g. NKG2D) or metabolic products derived from dysregulated tumors (e.g. mevalonate pathway). When activated, γδ T cells produce pro-inflammatory cytokines, such as interferon γ (IFNγ), which in turn promote anti-cancer CD8^+^ T cell immunity. They can also directly kill cancer cells via perforin/granzyme mediated cytotoxicity. Therefore, γδ T cell infiltration within the tumor bed is a favorable prognostic marker in several human cancers (7,8) and represents a promising target in cancer immunotherapy (9,10).

A growing body of evidence has also highlighted that the gut microbiota may have a positive impact on cancer prevention or treatment, especially via improved host immunosurveillance of cancer (11,12) as well as patients’ ability to respond to chemotherapy (13) or immunotherapy (11,14–16). These seminal studies have identified certain families, genera, or species of bacteria associated with the gut microbiota that appear to correlate with a strong host immune status and a subsequent favorable response to cancer therapy. For instance, *Bifidobacterium, Ruminoccoccaceae, Akkermansia muciniphila, Enterococcus irae*, and *Allistipes* have all been positively associated with ICB responses. As preclinical proof of concept, it has been shown that either Fecal Microbiota Transplantation (FMT) from responding patients or direct supplementation with immunostimulatory bacteria can restore the ability to respond to ICB in mice (14,15). Taken together, these studies strongly suggest that modulation of the gut microbiota may represent a major strategy to prevent and/or treat cancers by optimizing host immunity and subsequent anti-tumor responses.

It is noteworthy, 70% of host immune cells are located within the gut, close to microbiota-derived signals (17). Among the Intra Epithelial Lymphocytes (IELs) in the gut, the γδ T cells represent the largest immune subset. Interestingly, their early development is largely dependent on gut microbiota after birth (18). Unlike conventional αβ TcR^+^ CD4^+^ or CD8^+^ T cells, which recognize antigen-derived peptides, these cells recognize metabolic products derived from bacteria or tumor cells (19,20).

Diet is probably the most powerful microbiota modulator, in terms of both its composition and metabolic functions (21). Thus, modulation of the gut microbiota by nutritional intervention, particularly via prebiotics, could be of great interest in optimizing host immune responses, including cancer immunosurveillance. Prebiotics are defined as non-digestible food ingredients that benefit the host by selectively stimulating the growth and activity of one species or a limited number of bacterial species in the colon (22). Among the prebiotics, inulin, a Fructo-Oligo Saccharide derived from chicory roots, is well known to promote some colonic bacteria, including *Bifidobacterium*, which has been associated with optimal host immunity in the context of cancer immunosurveillance and immunotherapy (11,23). Recently, the use of inulin to stimulate immunosurveillance was confirmed by Li *et al*., who showed that an inulin diet attenuated tumor growth in a CD8^+^ T cell-dependent manner in mice (24). To date, the effect of nutritional intervention on systemic immunity, including the anti-tumor CD8^+^ T cell response, remains largely unknown.

Since an inulin-enriched diet is known to alter the metabolic activity of the gut microbiota (25), we hypothesized that intestinal and systemic γδ T cells might subsequently be altered. In the present study, we examined the role of γδ T cells in inulin-mediated anti-tumor effects and highlighted their pivotal role in CD8^+^ T cell–mediated immunosurveillance in several transplantable mouse tumor models.

## Results

### Inulin enriched diet leads to potent tumor growth control and enhanced γδ Tumor Infiltrated Lymphocytes activity

Inulin has been described as a promoter of immunostimulatory bacteria associated with strong anti-tumor immunity (11,26). Therefore, we evaluated whether 15 days of preconditioning in WT mice by adding inulin to drinking water could promote host anti-tumor immunity after subcutaneous injection of a syngeneic B16-OVA melanoma tumor (**Fig1A**). Consistent with a previous study in another tumor model (24), inulin strongly attenuated aggressive B16-OVA melanoma tumor growth (**Fig 1B left panel**). Notably, the inulin-mediated anti-tumor effect was also confirmed in other tumor models (i.e., MCA205 fibrosarcoma and MC38 colorectal cancer cell lines, **Fig 1B middle and right panel**). To better understand the mechanisms behind these observations, we analyzed the Tumor Infiltrating Lymphocytes (TILs) when the control diet group reached maximum ethical tumor size (day 13 post B16-OVA tumor inoculation, **Fig 1A**). Flow cytometry analyses revealed that inulin triggered greater infiltration of immune cells (CD45^+^ cells) within the tumor bed (**Fig 1C**) as well as greater tumor infiltration of dendritic cells (**Fig S1**). Consistent with the anti-tumor activity observed (**Fig 1B**), we found that the inulin-enriched diet triggered a potent Th1 anti-tumor response, as illustrated by the greater amount of IFNγ-producing CD4^+^ and CD8^+^ TILs (**Fig 1 D and E**). In agreement with this, immunization with OVA protein adjuvanted with PolyI:C (a Th1 polarizing adjuvant) led to a higher frequency of OVA-specific CD8^+^ T cells in the lymph nodes of inulin-supplemented mice (**Fig S2**). This suggests that inulin potentiated the Th1 polarizing activity of PolyI:C.

**Figure 1:**
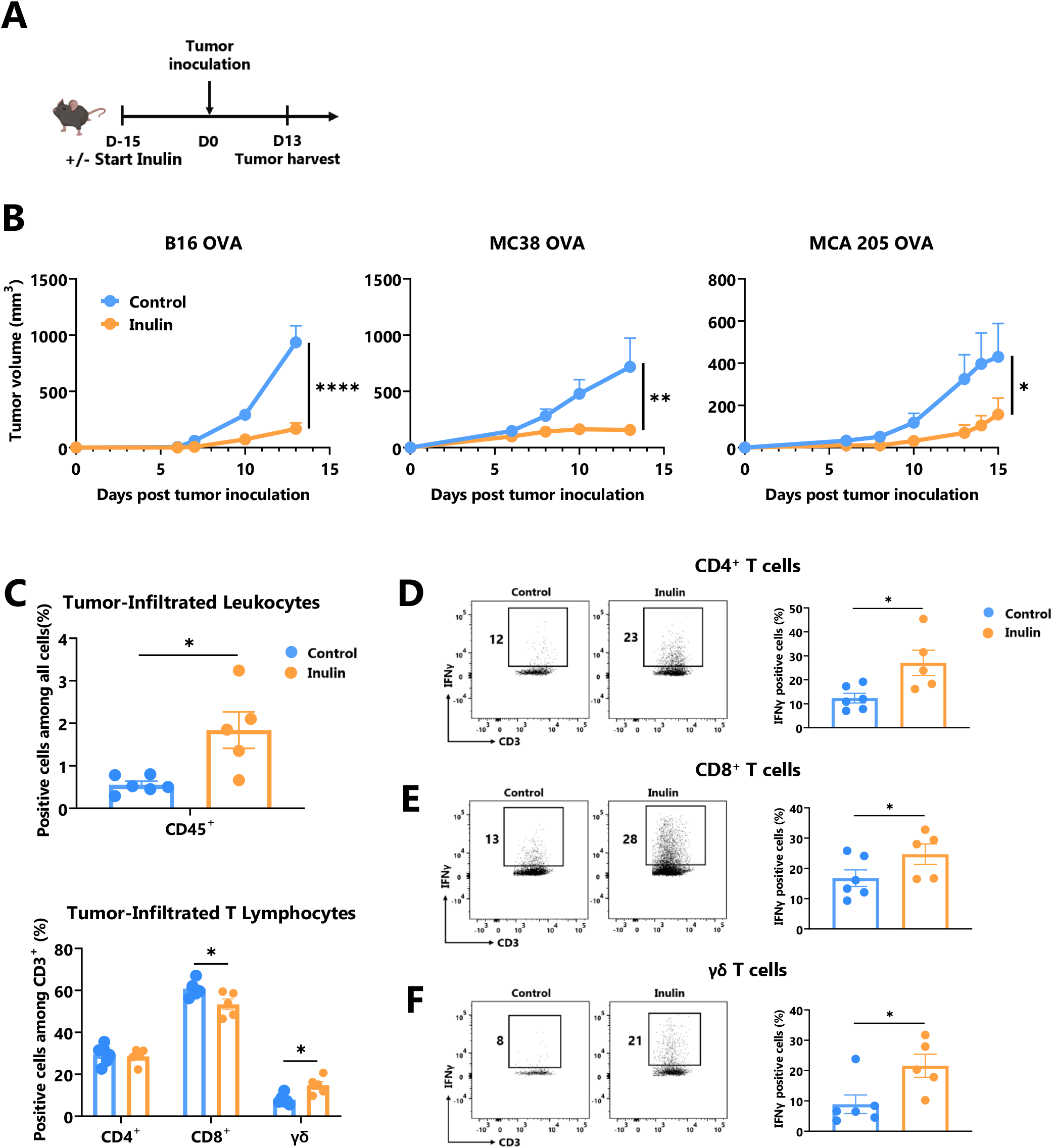
Inulin-enriched diet inhibits tumor growth and reinforces tumor infiltrated immunity. (A) Experimental schedule. C57BL/6 mice were fed with a control or an inulin-enriched diet (7.2% in drinking water) (n=6 mice per group) starting 15 days before subcutaneous (s.c.) inoculation of 2×10^5^ B16 OVA melanoma cells, or 5×10^5^ MC38 OVA colorectal cancer cells, or 2×10^5^ MCA 205 OVA fibrosarcoma cells (n=6 mice per group). (B) Tumor growth curves of mice treated as described in (A) and implanted with indicated tumors. (C) Frequency of B16 OVA tumor-infiltrated total CD45^+^ and effector CD4^+^, CD8^+^ or γδ TcR^+^ T Lymphocytes from mice treated as described in (A) when control tumors reached the maximal ethic size. (D-F) Frequency of B16 OVA tumor-infiltrated IFNγ-producing cells gated on CD45^+^ CD3^+^ (D) CD4^+^, (E) CD8^+^ or (F) γδ TcR^+^ from mice treated as in (A). Graphs show the mean ± SEM. Statistically significant results are indicated by: *p < 0.05, **p < 0.005, ****p < 0.0001 by two-way ANOVA with Geisser Greenhouse’s correction (B) or by Mann-Whitney tests (C-F).

In addition, we assessed the frequency and activity of γδ TILs, and found that inulin-enriched diet also promoted γδ TIL infiltration (**Fig 1C**) and IFNγ-production in residual tumors (**Fig 1F**). Notably, among both MCA205 and MC38 TILs, only γδ T cells infiltration was significantly increased under the inulin diet (**Fig S3**). Taken together, these data indicate that inulin triggered greater tumor infiltration by immune cells, potent Th1-polarized anti-tumor immunity, and in particular, high IFNγ production by conventional T cells as well as γδ TILs, which may ultimately lead to tumor growth control.

### Inulin-mediated anti-tumor immunity relies on γδ T cell

γδ T cells are unconventional T cells recognizing metabolic-related molecules such as phosphoantigens, and represent an interesting target/tool in cancer immunotherapy because they exhibit potent anti-tumor activity (6). As tumor infiltration and IFNγ production are enhanced by an inulin-enriched diet, we evaluated whether these cells could play a central role in inulin-mediated anti-tumor immunity. To assess their contribution, we used repeated injections of γδ TcR-blocking antibody starting the day before the tumor implantation (**Fig 2A**). Of note, while inulin-treated mice well-controlled tumor growth at day 13, after implantation, administration of γδ TcR-blocking antibody completely abrogated the inulin-mediated anti-tumor effect (**Fig 2B**). Interestingly, analysis of TILs revealed that under systemic γδ T cell blockade, the inulin-enriched diet failed to trigger greater IFNγ-production by CD4^+^ and CD8^+^ TILs (**Fig 2 C and D**). Furthermore, under γδ TcR blockade and despite the inulin diet, the frequency of IFNγ-producing γδ TILs was similar to that of the control group, suggesting that γδ TcR signaling is required in inulin-mediated immunoactivation (**Fig2E**). Taken together, these data indicate that γδ T cells, and most likely their TcR engagement, are mandatory for inulin-mediated anti-tumor effects. In addition, we observed a significant correlation between the level of tumor infiltration of IFNγ-producing γδ T cells and tumor growth in all three tumor models (**Fig 2F**), confirming their pivotal role in inulin-mediated anti-tumor effects.

**Figure 2:**
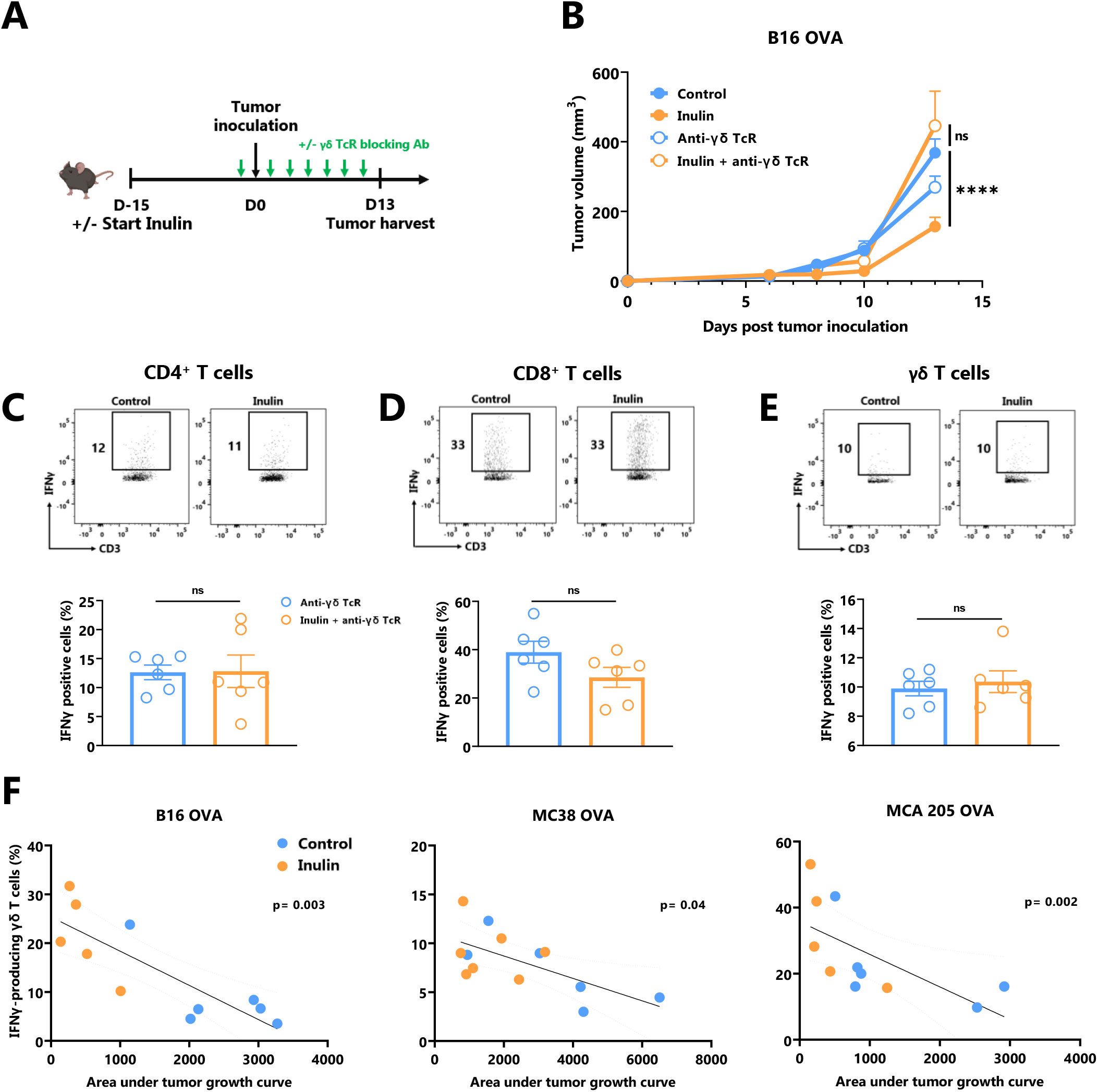
Inulin-triggered anti-tumor effect depends on γδ T lymphocytes. (A) Experimental schedule. C57BL/6 mice were fed with a control diet or inulin-enriched diet (7.2% in drinking water) (n=12 mice per group) starting 15 days before s.c. inoculation of 2×10^5^ B16 OVA melanoma cells. Anti-γδ TcR antibodies were injected intraperitoneally (i.p.) to 6 mice per group every two days starting the day before tumor inoculation. (B) Tumor growth curves of mice treated as described in (A). (C-E) Frequency of B16 OVA tumor-infiltrated IFNγ-producing cells gated on CD45^+^ CD3^+^ (C) CD4^+^, (D) CD8^+^ or (E) γδ TcR^+^ from mice treated as described in (A) when control tumors reached the maximal ethic size. (F) Correlation between IFNγ-producing γδ T cell infiltration in tumors and the tumor growth in mice fed on a control or inulin-enriched diet (n=6 mice per group), bearing the indicated tumor. (A-E) Graphs show the mean ± SEM, and (F) IFNγ-producing γδ T cell infiltration in the tumor depending to the tumor growth. ns = not-significant, ****p < 0.0001 by two-way ANOVA with Geisser greenhouse’s correction (B), by Mann-Whitney tests (C-E), or Spearman test (F).

### Inulin-enriched diet leads to gut-associated γδ T cell activation and gut inflammation

It is well known that the gut microbiota affects local (gut) and systemic immune tone (27). Since γδ T cells represent the most abundant immune cell population within the gut epithelium (28), we evaluated the impact of the inulin diet on IFNγ-production by gut-associated immune cells as well as on the level of gut epithelium inflammation. For this purpose, we collected intestines from mice following a 15-day inulin diet (**Fig 3A**). The fractions of Intra Epithelial Lymphocytes (IELs) and Lamina Propria Cells (LPCs) were analyzed using flow cytometry. Under inulin, while the frequency of IFNγ-producing CD4^+^ and CD8^+^ T cells was comparable to that of the control group in IELs and LPCs (**Fig 3B-C and Fig S4A, respectively**), γδ T cell IELs showed a higher proportion of IFNγ-producing cells (**Fig 3D**).

**Figure 3:**
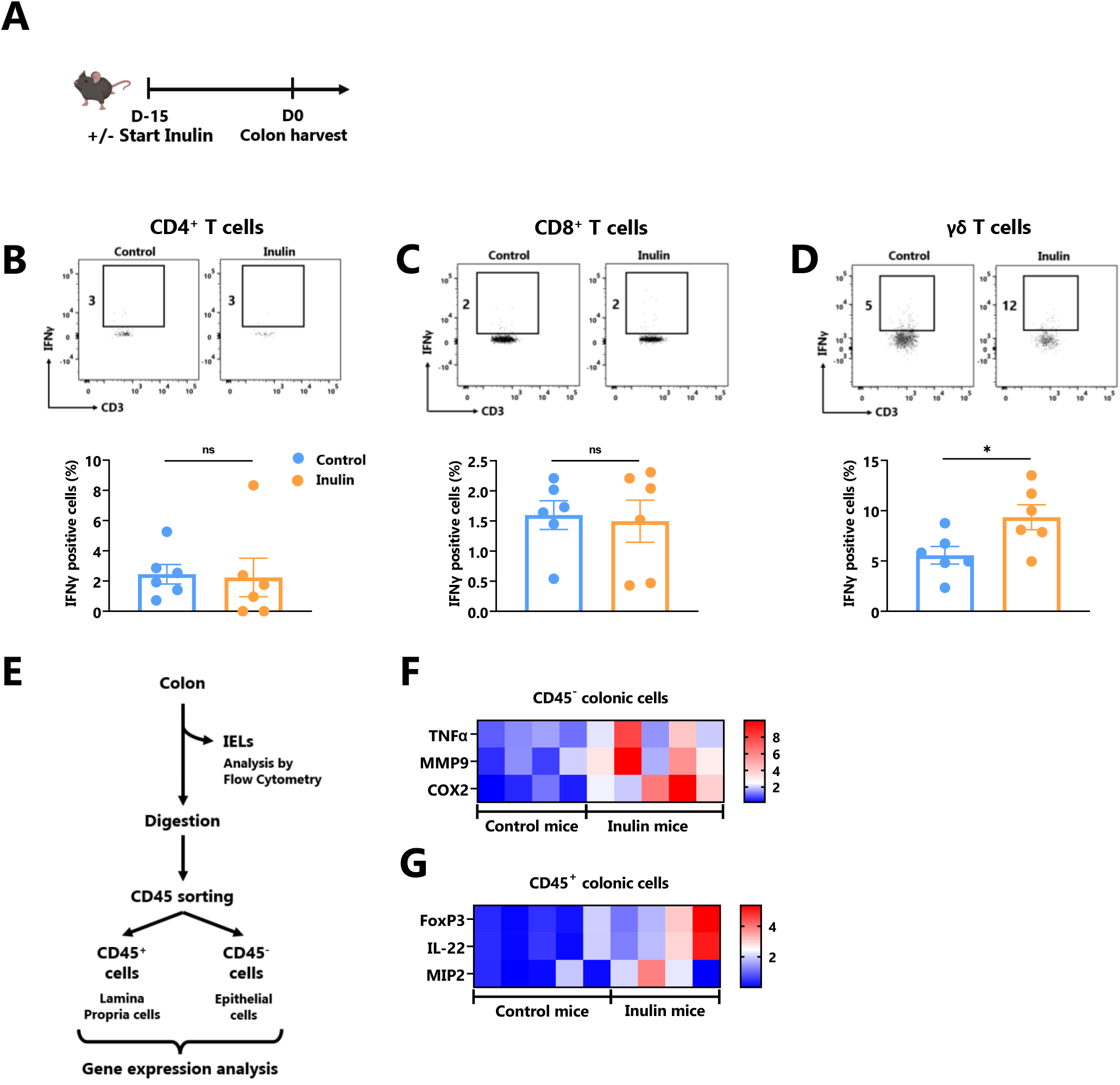
Inulin consumption promotes γδ T IELs and mucosa cell activation in the colon. (A) Experimental schedule. C57BL/6 mice were fed with a control diet or inulin-enriched diet (7.2% in drinking water) (n=12 mice per group) for 15 days before the analysis of their gut immunity. (B-D) Frequency of IFNγ-producing IntraEpithelial Lymphocytes (IELs) gated on CD45^+^ CD3^+^ (B) CD4^+^, (C) CD8^+^, or (D) γδ TcR^+^ from mice treated as described in (A). (E) Experimental plan of colon treatment after harvest. After isolation of IELs, remaining colon pieces were digested before sorting of CD45^-^ and CD45^+^ cells for further analysis. (F-G) qRT-PCR analysis of inflammation, tissue repair and tight junction -related genes in (F) CD45^-^ and (G) CD45^+^ colon cells sorted as described in (E). (B-D) Graphs show the mean ± SEM. ns = not-significant, *p < 0.05 by Mann-Whitney tests (B-D). (F-G) Graphs show the expression levels of significantly increased genes (p < 0.05 by Mann Whitney test).

To further characterize inulin-mediated intestinal inflammation, we sorted the epithelial (CD45-) fraction from LPCs (CD45^+^) to perform RT-qPCR on genes involved in inflammation, tissue repair, and tight junctions (**Fig 3 E-G, Fig S5, and Table S2**). Of the 19 analyzed genes, three inflammation-related genes were significantly upregulated under the inulin regime in epithelial cells, namely Tumor Necrosis Factor (TNF)-α, Cyclo-Oxygenase (Cox)-2 and Matrix Metalloprotease (MMP)-9 (**Fig 3F**). In the CD45^+^ compartment (LPCs), inulin triggered significant overexpression of Macrophage Inflammatory Protein (MIP)-2, Interleukin (IL)-22 and the transcriptional factor Forkhead Box P (FoxP)-3 (**Fig 3G**). Similarly, we observed a trend toward higher IL-10 production, although not significant, by LPC-associated T cells (**Fig S4B**). Altogether, these results indicate that inulin diet triggered both targeted γδ T cell activation in IELs and inflammation in gut epithelial cells, in parallel with tissue repair signaling and tolerance induction in the *lamina propria*.

### *Inulin-enriched diet alters the gut microbiota and promotes* Bifidobacterium *growth*

To establish a link between the observed intestinal immune activation (**Fig 3**) and the inulin-enriched diet, we characterized the composition of fecal bacterial microbiota after inulin administration using 16S rDNA sequencing. Metagenomic analysis indicated that five phyla composed the microbiota of these mice: *Firmicutes* (58% of the sequences), *Bacteroidota* (37% of the sequences), *Desulfobacterota* (1.3% of the sequences), *Proteobacteria* (1.2% of the sequences), and *Actinobacteriota* (2.4% of the sequences). A 15-day inulin diet profoundly altered the gut microbiota composition, as revealed by the relative abundance of these bacterial phyla observed in inulin-treated mice compared to control mice (**Fig 4A**), with a visible increase in bacteria of the *Actinobacteriota* phylum, and more precisely to the genus *Bifidobacterium*. This diet altered the alpha diversity; the total number of species was not significantly different (387±53 Operational Taxonomic Units (OTUs) for the control group and 378±43 for the inulin group), but the distribution of sequences within the OTUs was significantly different (Shannon index, p = 0.00582). A Bray-Curtis β-diversity analysis illustrated by Principal Component Analysis (PCoA) representation confirmed a significant clustering of the overall bacterial microbiota with a p-value < 0.001 (**Fig 4B**). To identify whether particular bacteria were potentially involved in anti-tumor immunity, we performed differential abundance correlation using a linear discriminant analysis (LDA) effect size algorithm (LEfSe; see Material and Methods). The LDA scores plotted in a phylogenic cladogram showed significant differences across several taxonomic ranks (**Fig 4C**). One of the strongest signals, LDA score>4, was the increase in *Bifidobacterium animalis* subsp. *animalis*, a member of the *Actinobacteriota* phylum, in inulin treated mice (4.67%), compared with control mice (0.03%) (**Fig4D, Table S1**). Importantly, consistent with our results, *Bifidobacterium* has been shown to exhibit immunostimulatory properties in the context of cancer immunosurveillance and immunotherapy (11,23). Therefore, *Bifidobacterium* may be involved in the stimulation of inulin-mediated anti-tumor immunity. The second most significant signal concerned the family Clostridiaceae, and in particular, the genus *Clostridium sensu stricto 1*. Our sequences were close to *those of Clostridium saudiense, Clostridium disporicum*, and *Clostridium celatum* (these species have nearly identical 16S rDNA sequences). This OTU statistically decreased in the mice microbiota due to treatment, from 6% in the control group to 0,09% in the inulin group (data not shown).

**Figure 4:**
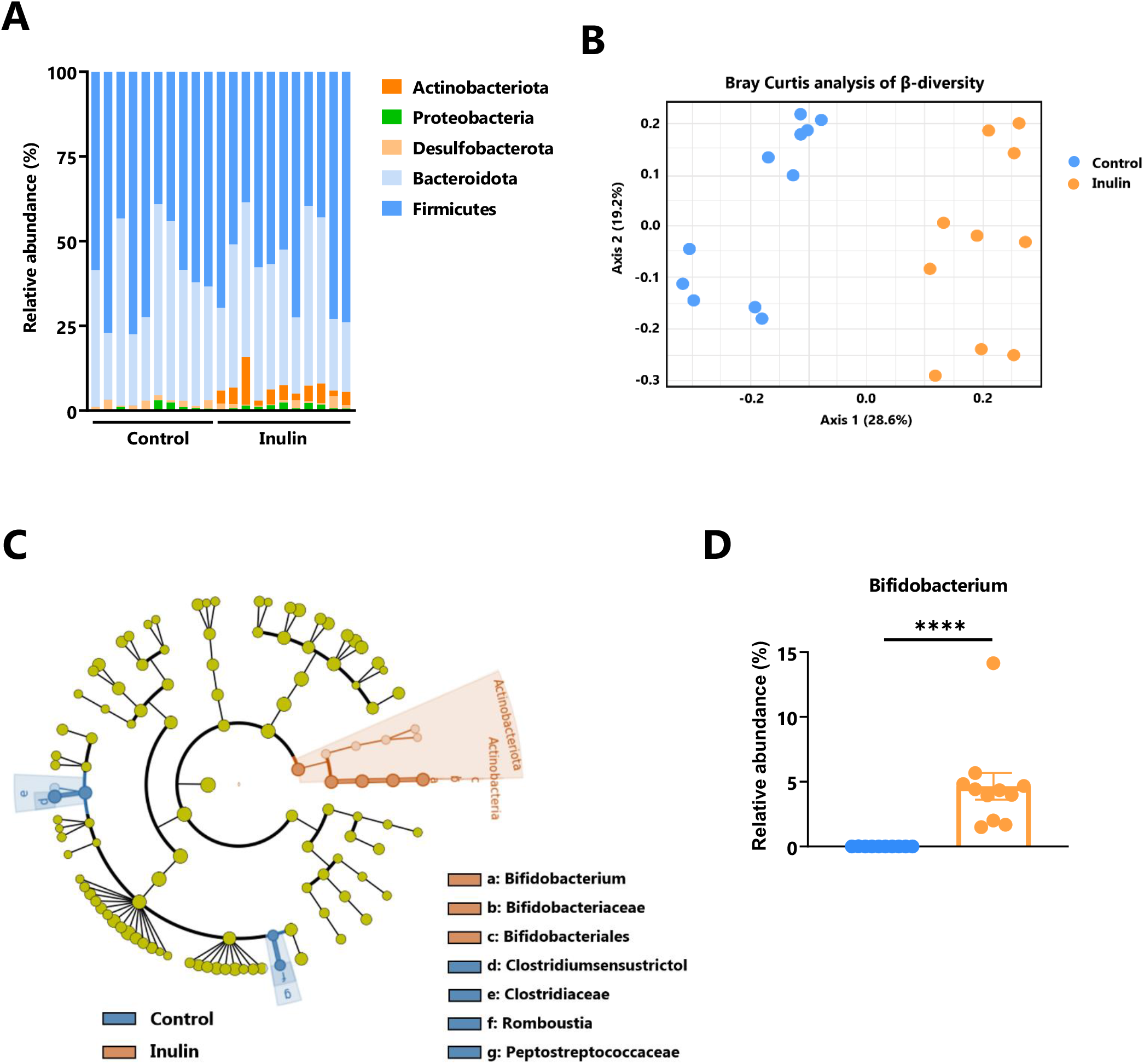
Inulin modulates gut microbiota and promotes *Bifidobacterium* growth. Feces from C57BL/6 mice fed with a control (n=10 mice) or an inulin-enriched diet (7.2% in drinking water) (n=11 mice) for 15 days were collected and analysed by 16S rDNA sequencing. (A) Relative abundance of bacterial phyla (B) Bray-Curtis analysis of β-diversity, (C) cladogram representation of Linear discriminant analysis Effect Size (LEfSe) analysis, and (D) relative abundance of Bifidobacterium genus. The graph shows the mean ± SEM. ****p < 0.00001 by by Mann-Whitney test.

### Inulin-mediated increased anti-tumor immunity relies on gut microbiota

Inulin, a polysaccharide molecule, has been shown to possess adjuvant properties *in vivo* (29,30). Such direct immune activation may be responsible for the anti-tumor immunity observed under the inulin regime. To address this issue, we administered a broad-spectrum antibiotic cocktail to the mice 2 days before the inulin diet to significantly reduce the number of bacteria in their gut, as previously described (31) (**Fig 5A**). After tumor implantation, we observed that concomitant antibiotic treatment completely abolished the ability of inulin to attenuate tumor growth (**Fig 5B**). Furthermore, inulin failed to trigger a higher frequency of IFNγ-producing γδ TILs and subsequent CD4^+^ and CD8^+^ TILs (**Fig 5 C, D, and E**). Altogether, these findings indicate that inulin-mediated anti-tumor activity cannot be attributed to a direct adjuvant effect; rather, it relies on host immune activation mediated by the gut microbiota.

**Figure 5:**
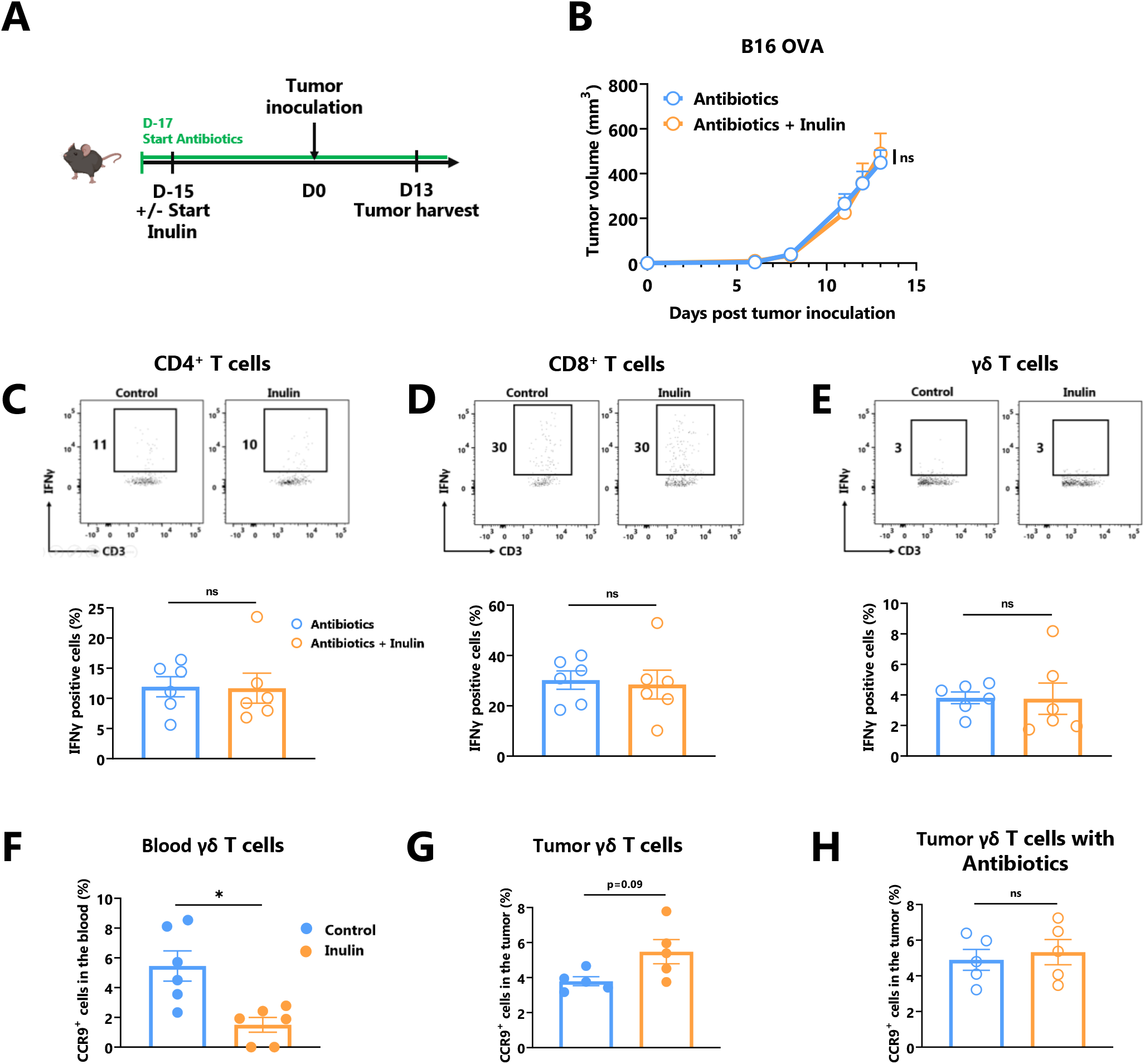
Inulin-triggered anti-tumor effect relies on intact bacterial gut microbiota. (A) Experimental schedule. C57BL/6 mice were fed with a control diet or inulin-enriched diet (7.2% in drinking water) (n=6 mice per group) starting 15 days before s.c. inoculation of 2×10^5^ B16 OVA melanoma cells. Microbial gut microbiota was depleted from 2 days before the beginning of the diet by adding vancomycin (0.25 g/L), colimycin (12×10^6^ U/L), ampicillin (1 g/L) and streptomycin (5 g/L) to drinking water. (B) Tumor growth curves of mice treated as described in (A). (C-E) Frequency of B16 OVA tumor-infiltrated IFNγ-producing cells gated on CD45^+^ CD3^+^ (C) CD4^+^, (D) CD8^+^, or (E) γδ TcR^+^ from mice treated as described in (A). (F-G) Frequency of CCR9^+^ cells among CD45^+^ CD3^+^ γδ TcR^+^ in (F) the blood and in (G) the tumor of mice fed with a control diet or inulin-enriched diet (7.2% in drinking water) (n=6 mice per group) starting 15 days before s.c. inoculation of B16 OVA melanoma cells. (H) Frequency of CCR9^+^ cells among CD45^+^ CD3^+^ γδ TcR^+^ in tumor of mice treated as described in (A). Graphs show the mean ± SEM. ns = not-significant, *p < 0.05 by two-way ANOVA with Geisser greenhouse’s correction (B) or by Mann-Whitney tests (C-H).

Because γδ T cells are obligatory for inulin-mediated anti-tumor immunity (**Fig 2**), and that γδ T IELs are the only T cell subset activated by inulin diet in the gut (**Fig 3**), we hypothesized that γδ TILs could have an intestinal origin. To address this question, we analyzed the expression of the gut-homing chemokine receptor CCR9 in circulating γδ T and γδ TILs. Whereas the number of CCR9-expressing γδ T decreased in the blood (**Fig 5F**), their frequency tended to increase in the tumor bed, despite a possible loss of expression in this environment (**Fig 5G**). Notably, under a concomitant inulin diet and antibiotic treatment, γδ TILs expressed similar levels of CCR9 compared with the control group (**Fig 5H**). Taken together, these data suggest that inulin-induced activated γδ TILs may be of intestinal origin.

### Inulin regimen is as efficient as αPD-1 Immunotherapy

A large body of evidence suggests that certain immunostimulatory bacteria, including *Bifidobacterium*, may facilitate the efficacy of immune checkpoint inhibitors, including those targeting the anti-PD1/PD-L1 axis (11,14,15,32). Because the inulin diet altered the gut microbiota (**Fig5**) and favored *Bifidobacterium* (**Fig 4**), we evaluated whether this diet could be synergistic with anti-PD1 treatment (**Fig 6A**). Tumor growth monitoring indicated that inulin alone allowed tumor growth control to the same extent as anti-PD-1 alone or in combination (**Fig 6B**). The individual mouse curves clearly showed that inulin alone triggered the strongest anti-tumor effect compared with anti-PD1 alone or in combination (**Fig 6C**). TILs analysis revealed that inulin led to a greater IFNγ production by γδ T cells (**Fig 6D**) and CD4^+^ cells (**Fig 6E**), compared with anti-PD1 alone. The combination did not potentiate this effect despite similar PD-1 expression among the TILs (**Fig S6**). Anti-PD1 immunotherapy reinvigorates CD8^+^ TILs and increases their ability to produce IFNγ (33). In agreement with this, anti-PD1 treatment led to a higher proportion of IFNγ-producing CD8^+^ TILs within the tumor bed (**Fig 6F**). Interestingly, inulin alone triggered the activation of CD8^+^ TILs, at least to a similar extent.

**Figure 6:**
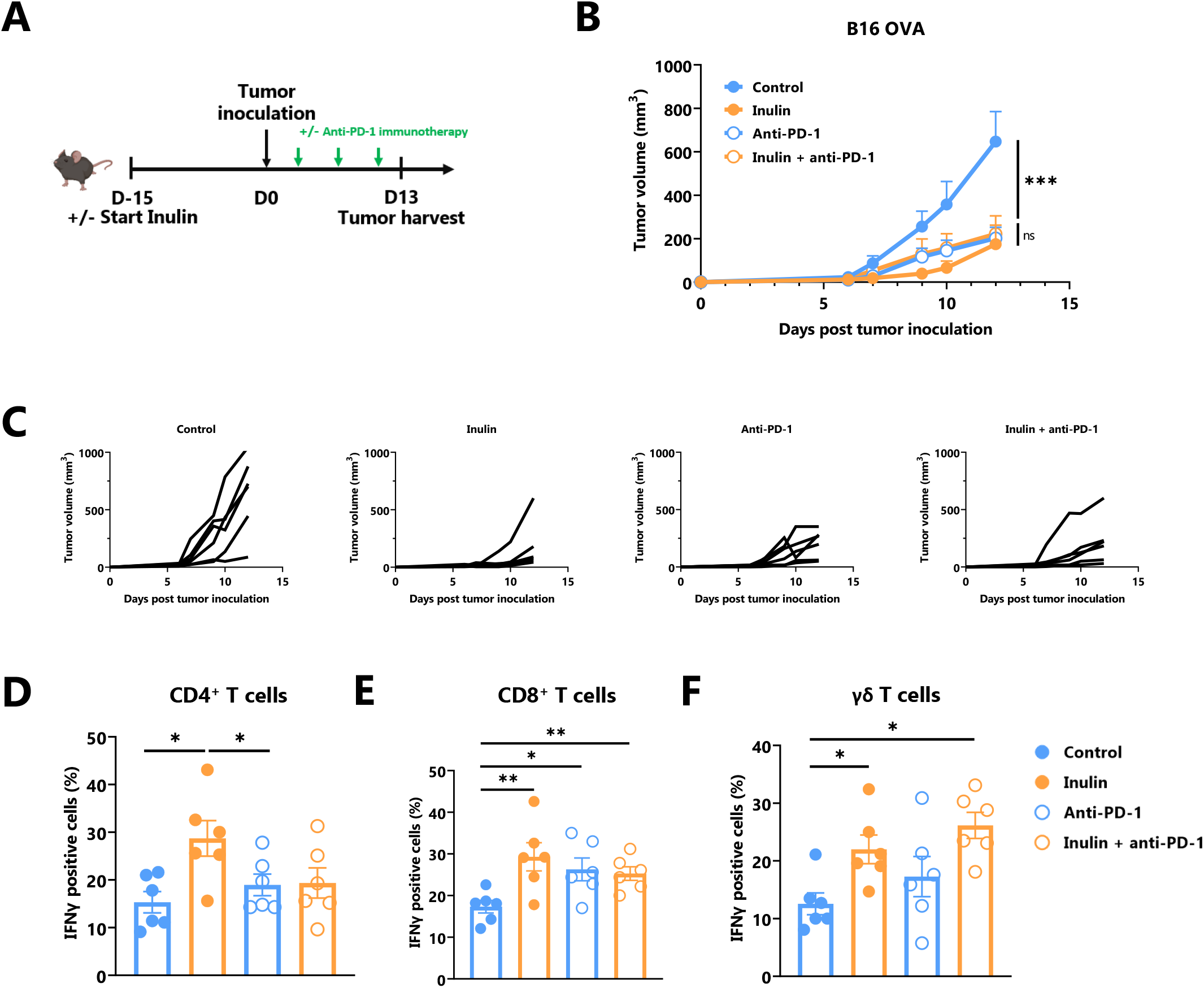
Inulin treatment is as efficient as anti-PD-1 immunotherapy. (A) Experimental schedule. C57BL/6 mice were fed with a control diet or inulin-enriched diet (7.2% in drinking water) (n=12 mice per group) starting 15 days before s.c. inoculation of 2×10^5^ B16 OVA melanoma cells. Anti-PD-1 immunotherapy was injected i.p. to 6 mice per group on days 4, 7 and 10 post-tumor inoculation. (B) Mean and (C) individual tumor growth curves of mice treated as described in (A). (D-F) Frequency of B16 OVA tumor-infiltrated IFNγ-producing cells gated on CD45^+^ CD3^+^(C) CD4^+^, (D) CD8^+^, or (E) γδ TcR^+^ from mice treated as described in (A). Graphs show the mean ± SEM. ns = not-significant, *p < 0.05, **p < 0.005, ***p < 0.001 by two-way ANOVA with Geisser greenhouse’s correction (B) or by Mann-Whitney tests (D-F).

## Discussion

Gut microbiota is emerging as a powerful host immune modulator that can be exploited for both cancer prevention and treatment. Consequently, there is an urgent need to understand the cellular and molecular mechanisms that govern this modulation to rapidly translate these findings into innovative preventive or therapeutic strategies.

Prebiotics contribute to the modification of the composition and functions of the microbiota, and subsequently, to the modification of host immunity. The importance of a prebiotic-enriched diet was recently highlighted in humans by Spencer *et al*., who reported that a fiber-enriched diet positively influenced the response to immunotherapy in melanoma patients (34).

Inulin is a widely consumed prebiotic, either in the diet or as a supplement. Recently, inulin was reported to attenuate transplantable melanoma tumor growth (24) in a CD8^+^ T cell-dependent manner. In the current study, we reported that inulin-enriched diet triggers not only potent αβ T cell anti-tumor immunity, but also the accumulation and activation of intratumoral γδ T cell in a microbiota-dependent manner. We emphasized that these cells are indispensable for the increase of IFNγ-producing CD4^+^ and CD8^+^ TILs and subsequent tumor growth control. We also reported that an inulin-enriched diet remarkably alters the composition of the microbiota and, as expected, significantly promotes the selective growth of *Bifidobacterium* species, which are known to be immunostimulatory. Consistent with this, such a diet promotes the activation of γδ T IELs in the gut and we found that the frequency of CCR9^+^ tumor-infiltrating cells tends to increase, suggesting a possible intestinal origin of γδ TILs. Interestingly, CCR9-expressing T cells have been shown to contribute to anti-tumor immunity, and their recruitment into the tumor-bed (by CCL25 intratumoral delivery) (35) improves immunotherapy efficacy.

Regarding a possible synergy with immunotherapy, we found that a prophylactic inulin diet was as effective as anti-PD-1 immunotherapy in our setting and that PD-1 expression was equivalent between groups. Because PD-1 expression identifies exhausted T cells only when combined with other inhibitory receptors such as Tim-3, LAG-3, or TIGIT (36), it is possible that PD-1^+^ TILs under inulin were not in an exhausted state, in contrast to TILs in the control group. Furthermore, in our experiment, we only harvested the tumors 13 days after inoculation to characterize the functionality of the TILs, so it is possible that the combination of inulin and anti-PD-1 treatment would prevent later tumor escape compared to anti-PD-1 alone in the B16 melanoma model. To fully address this question, further phenotyping of TILs and prolonged monitoring of tumor growth are required.

To date, how intestinal and/or systemic γδ T cells are activated by an inulin-enriched diet remains unclear. Importantly, we observed that γδ TcR blockade abrogated inulin-mediated enhanced immunosurveillance. Beyond demonstrating that γδ T cells are obligate mediators, these data highlighted that their activation was TcR-dependent and thus that microbiota-derived γδ T cell activation signals could be metabolite-mediated. Altogether, these data allow us to propose a scenario in which intestinal γδ T cells could be activated in a TcR-dependent manner, most likely by metabolites derived from inulin-mediated remodeling of the microbiota and then recirculated to the tumor to promote CD8^+^ T cell anti-tumor activity.

In this study, we report three transplantable tumor models in which inulin triggered anti-tumor immune responses. As a result, the growth of melanoma, colorectal and fibrosarcoma tumors were attenuated and accompanied by an increased in γδ T cell infiltration. To fully investigate the impact of nutritional intervention on cancer immunosurveillance mediated by γδ T cells, the use of more physiological models, such as genetically engineered mouse models or chemically induced cutaneous tumor models (4), may be useful as they allow for the establishment of proper immunoediting, unlike transplantable tumors.

Despite these limitations, our data provide a proof of concept that intestinal and systemic γδ T cell functions can be modulated by nutritional intervention. In particular, we identified these cells as a critical immune subset that is mandatory for inulin-mediated anti-tumor immunity and subsequent control of tumor growth *in vivo*. These results further support and rationalize the use of such prebiotic approaches as well as the development of immunotherapies targeting γδ T cells in cancer prevention and immunotherapy. Notably, determining which bacteria-derived metabolite(s) is responsible for this activation would be of great interest in developing future postbiotic immunotherapeutic approaches and rapidly translating these findings for cancer prevention and treatment in humans.

## Acknowledgements

We would like to thank Sylvie Berthier (Cytometry Platform, CHUGA), Dr. Hervé Lerat, Kevin Escot and Laurie Arnaud (PHTA Animal Facility), Clément Caffaratti and Hélène Coradin (TIMC Lab), and Théo Ziegelmeyer (Inserm U1209) for their technical assistance.

This work (D.H.) was supported by GEFLUC Dauphiné-Savoie, Ligue contre le Cancer Comité Isère, Ligue contre le Cancer Comité Savoie, Université Grenoble Alpes IDEX Initiatives de Recherche Stratégiques. E.B. is supported by a grant salary from the French Ministry of Higher Education, Research and Innovation. The funders had no role in study design, data collection or analysis.

This work benefited from the facilities and expertise of @BRIDGe for 16S sequencing (Université Paris-Saclay, INRAE, AgroParisTech, GABI, 78350 Jouy-en-Josas, France).

## Author’s contribution

Conceptualization: DH, EB, BT

Performed experiment: DH, EB, CP

Analyzed data: DH, EB, CP, MLR, IM, AS

Wrote the paper: DH

Contribution to paper edition/writing: EB, CP, BT, DA, MC, MLR, IM, and AS

Grants acquisition: DH

## Material and Methods

### Animals

Female C57Bl/6 mice (5 to 6 weeks aged) were provided by Janvier SA Laboratory (Le Genest-Saint-Isle, France) and housed at “Plateforme de Haute Technologie Animale (PHTA)” UGA core facility (Grenoble, France), EU0197, Agreement C38-51610006, under specific pathogen-free conditions, temperature-controlled environment with a 12-h light/dark cycle and ad libitum access to water and diet. Animal housing and procedures were conducted in accordance with the recommendations from the Direction des Services Vétérinaires, Ministry of Agriculture of France, according to European Communities Council Directive 2010/63/EU and according to recommendations for health monitoring from the Federation of European Laboratory Animal Science Associations. Protocols involving animals were reviewed by the local ethic committee “Comité d’Ethique pour l’Expérimentation Animale no.#12, Cometh-Grenoble” and approved by the Ministry of Research (APAFIS#12905-2018010411002729-v5). Mice were distributed in groups according to their diet. For inulin treatment, mice received standard diet and drinking water supplemented with 7.2% inulin starting 15 days before tumor inoculation. Drinking bottles supplemented with inulin were renewed 3 times a week.

### Cell lines

Ovalbumin-expressing B16 melanoma (B16-OVA), MCA 205 fibrosarcoma cell lines (MCA 205 OVA), and MC38 colorectal cancer cell line (MC38 OVA) were kindly provided by Pr. L. Zitvogel, Gustave Roussy (B16-OVA, MCA205-OVA) and C. Fournier, Inserm U1209 (MC38-OVA). B16-OVA and MC38-OVA cell lines were cultured in 10% Fetal Bovine Serum (FBS) (Gibco) Dulbecco’s modified Eagle’s medium (DMEM) (Gibco) complete medium (supplemented with 1% non-essential amino acids, 1 mM sodium pyruvate, 50 U/mL penicillin, and 50 μg/mL streptomycin (all from Gibco)). MCA 205 OVA cell line was cultured in 10% FBS Roswell Park Memorial Institute (RPMI) complete medium (Gibco). For plasmid selection, G418 (500μg/ml, Sigma) and hygromycin B (50μg/ml, Sigma) antibiotics were added to B16-OVA/MC38-OVA and MCA 205-OVA cultures, respectively. All cell lines were tested as mycoplasma-free.

### Tumor implantation

Mice received subcutaneous implantation of either2 × 10^5^ B16-OVA cells, 2 × 10^5^ MCA 205-OVA cells, or 5 × 10^5^ MC38-OVA cells in 100 μL PBS in the right flank. Once palpable, tumors were measured using a caliper, and tumor volume was determined using the following formula: V_tumor_ = 0.5 x (width x lenght^2^).

### In vivo injection of anti-γδ TCR and anti-PD-1 antibodies

Mice received 100 μg of the monoclonal antibody anti-γδ TcR (clone UC7-13D5, Bio X Cell) or anti-PD-1 (clone RMP1-14, Bio X Cell) intraperitoneally, in PBS. Anti-γδ TcR antibodies were injected the day before tumor implantation and then every 2 days until the end of the experiment. Anti-PD-1 immunotherapy was administered on days 4, 7, and 10 after tumor implantation.

### In vivo bacterial microbiota depletion

Mouse gut microbiota was depleted using a cocktail of antibiotics diluted in drinking water: Vancomycin 0.25 g/L (Mylan), Colimycin 12 × 10^6^ U/L (Medac), Ampicillin 1 g/L (Sigma) and Streptomycin 5 g/L (Sigma), as previously described (13).

### Tumor digestion

Tumors were collected in complete RPMI medium, lacerated with scalpels, and digested with Liberase™ 2.5 mg/mL (Roche). Finally, tumors were crushed through a 70 μm cell strainer, washed, and resuspended in 10% FBS complete RPMI.

### Colonic IELs and LPCs isolation and CD45^+/-^ cell sorting

Colons were collected in Hank’s Balanced Salt Solution without Ca^2+^ and Mg^2+^ (HBSS^-^) (Gibco), then digested twice with 5 mM HEPES, 5 mM EDTA, 1 mM DTT, and 5% FBS, for IELs isolation. remaining colon pieces were then lacerated with scalpels and digested with collagenase VIII 0.5 mg/mL, DNAse I 10 U/mL (Sigma), and 5% FBS in complete RPMI for LPCs isolation. Finally, digested colon pieces were crushed through a 70 μm cell strainer, and cells were resuspended in 10% FBS complete RPMI. CD45^+^ and CD45^-^ cells from the LPCs cell fraction were sorted using MACS (Miltenyi Biotec) with CD45 microbeads, according to the manufacturer’s protocol.

### Lymph node digestion

Lymph nodes were collected in complete RPMI and then crushed through a 70 μm cell strainer. Cells were later suspended in 10% FBS complete RPMI.

### Spleen digestion

Spleens were collected in complete RPMI, then crushed using a 70 μm cell strainer, and washed with 10% FBS RPMI. The pellets were then suspended in 500 μL of Red Blood Cell Lysis buffer 1X (Sigma) and washed with 10% FBS complete RPMI.

### Blood sample preparation

Blood was collected via retro-orbital sampling in K2E tubes (BD Medical). After centrifugation, blood pellets were resuspended in 1 mL Red Blood Cell Lysis buffer 1X (Sigma) and washed with 10% FBS complete RPMI.

### Flow cytometry

To allow intracellular labelling of cytokines, cell suspensions were stimulated 4 h at 37°C with 50 ng/mL phorbol 12-myristate 13-acetate (PMA) (Sigma), 1 μg/mL ionomycin (Sigma), and Golgi Stop™ (BD Biosciences). For extracellular labelling, cells were incubated with 200 ng of each antibody. Antibodies targeting extracellular proteins were CD45 (30-F11), PD-1 (RMP1-30) (BD Biosciences), CD3 (17A2), CD4 (GK1.5), CD44 (IM7), CCR9 (9B1), CD11c (N418), CD8a (53-6.7), PD-L2 (24F.10C12), PD-L1 (10F.9G2), MHC-class II (M5/114.5.2), CD80 (16-10A1), CD40 (3/23) (Biolegend), CD11b (M1/70), γδ-TcR (eBioGL3 (GL-3, GL3), and OVA-dextramer (H-2 Kb) (Immudex) (**Table S2**).

To allow intracellular labelling, cells were first permeabilized using a FoxP3 staining buffer kit (eBioscience). Antibodies targeting intracellular cytokines were IFNγ (XMG1.2) (Biolegend), IL-10 (JES5-16E3) (Thermo Fisher Scientific), and IL-17A (TC11-18H10) (BD Biosciences) (**Table S2**). After intracellular labelling, cells were fixed and stored using FACS Lysing Solution 1X (BD Biosciences). All data were collected using a BD Biosciences FACSCanto II or Lyric and analyzed using FlowJo software.

### Bacterial DNA extraction and 16S sequencing from mouse feces

Fecal samples were collected and stored at -80°C. DNA was extracted from feces following optimized JJ Godon protocole (37). After nucleic acid precipitation with isopropanol, DNA suspensions were incubated overnight at 4°C and centrifuged at 20,000 × g for 30 min. The supernatants were transferred to a new tube containing 2 μL of RNase (RNase A, 10 mg/ml; EN0531; Fermentas, Villebon sur Yvette, France) and incubated at 37°C for 30 min. Nucleic acids were precipitated by the addition of 1 ml of absolute ethanol and 50 μl of 3 M sodium acetate and centrifuged at 20,000 × g for 10 min. The DNA pellets were washed with 70% ethanol 3 times, dried, and resuspended in 100 μl of Tris-EDTA (TE) buffer (10 mM Tris-HCl, 1 mM EDTA, adjusted pH 8). The DNA suspensions were stored at -20°C for 16S rDNA sequence analysis.

### 16S sequencing data processing

Bacterial diversity was determined for each sample by targeting a portion of the ribosomal genes. PCR was performed to prepare amplicons using V3-V4 oligonucleotides (PCR1F_460: 5’ CTTTCCCTACACGACGCTCTTCCGATCTACGGRAGGCAGCAG 3’, PCR1R_460: 5’ GGAGTTCAGACGTGTGCTCTTCCGATCTTACCAGGGTATCTAATCCT 3’), with 30 amplification cycles at an annealing temperature of 65°C. Amplicon quality was verified by gel electrophoresis, and PCR products were sent to the @BRIDGe platform for sequencing on an Illumina MiSeq platform (Illumina, San Diego, CA, USA).

The 16S sequences were demultiplexed and quality-filtered using the QIIME version 2.1.0 software package (38).. Sequences were analyzed and normalized using the pipeline FROGS (Find Rapidly OTU with Galaxy Solution) (39,40). PCR primers were removed and sequences with sequencing errors in the primers were excluded. The reads were clustered into Operational Taxonomic Units (OTUs) using the Swarm clustering method. Chimerae were removed, and 580 OTUs were assigned at different taxonomic levels (from phylum to species) using the RDP classifier and NCBI Blast+ on the Silva_138.1_16S database. When needed, the phylogeny was checked using the Phylogeny browser (https://www.ncbi.nlm.nih.gov/Taxonomy/Browser/wwwtax.cgi).

35538 reads were randomly selected from each sample to normalize the data. Raw reads were deposited into the SRA database (ID: PRJNA888063). The R packages “Biom” and “Phyloseq” were used for data analysis and plotting (41). Rarefaction analysis was performed to compare the relative abundance of OTUs across samples. Alpha diversity was measured using Observed, Chao1, Shannon, Simpson, and Inverse Simpson indexes. Beta diversity was measured using the Bray-Curtis distance matrix and used to construct principal coordinate analysis (PCoA) plots. The linear discriminant analysis (LDA) effect size (LEfSe) algorithm was used to identify taxa specific to diet and/or treatment (39,40).

### Real-time quantitative Reverse Transcription (qRT PCR)

Total RNA was extracted from IELs and LPCs using the RNEASY-QIAGEN kit, according to the manufacturer’s protocol. The purity and concentration of the extracted RNA were determined by reading the absorbance at 260 and 280 nm using a NanoDrop spectrophotometer (Thermo Fisher). The extracted RNA was then retrotranscribpted using the Superscript II kit (Invitrogen). qRT-PCR analysis was performed using SYBR Green Master Mix (Applied Biosystems) and the StepOne™ RealTime PCR system (Thermo Fischer). PCR conditions were: 2 min at 95°C, then 40 cycles of 15 s at 95°C and 1 min at 60°C. The sequence-specific primers used are shown in **Table S3** (GAPDH and β2M primers were from Eurofins Scientific; other primers were from Merck). The expression levels were normalized to GAPDH and β2M endogenous gene levels, and each value was then compared to a reference sample named the calibrator. Finally, relative quantification (RQ) values were calculated.

