## Supplemental data and Material and Method for "Inulin prebiotic reinforces host cancer immunosurveillance via γδ T cell activation"

### Supplementary Data and Tables

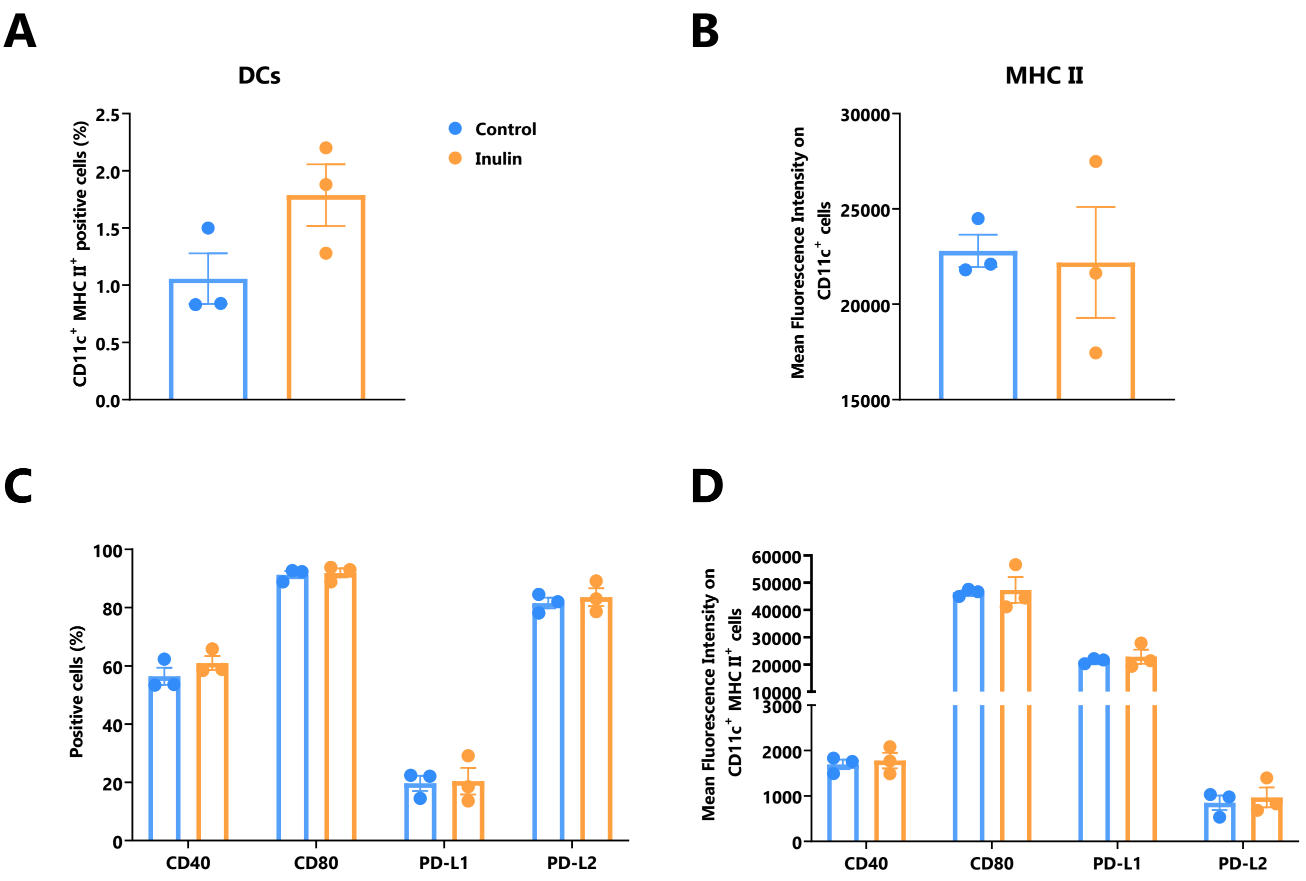


**Supplemental Figure S1: Inulin-enriched regimen leads to higher DC infiltration in B16 OVA tumors.**

(A) Frequency of Dendritic Cells (DCs) among CD45$^{+}$ cells in the B16 OVA tumor of mice treated as described in Figure 1A. (C) Frequency and (B-D) MFI of DC activation markers on DCs in the B16 OVA tumor of mice treated as described in Figure 1A. Graphs show the mean ± SEM.

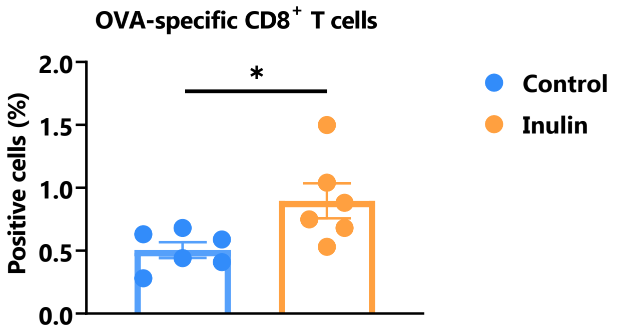


**Supplemental Figure S2: Inulin potentiates antigen-specific CD8+ T cell response after OVA PolyI:C immunization.**

C57BL/6 mice were fed with a control diet or inulin-enriched diet (7.2% in drinking water) (n=12 mice per group) starting 15 days before s.c. immunization against OVA protein (500µg/mouse) adjuvantized with PolyI:C (50µg/mouse). Frequency of OVA-specific CD8+ T cells in vaccine draining (inguinal) lymph nodes was analyzed 7 days post-immunization. Graph shows the mean ± SEM. *p < 0.05, by Mann-Whitney test.

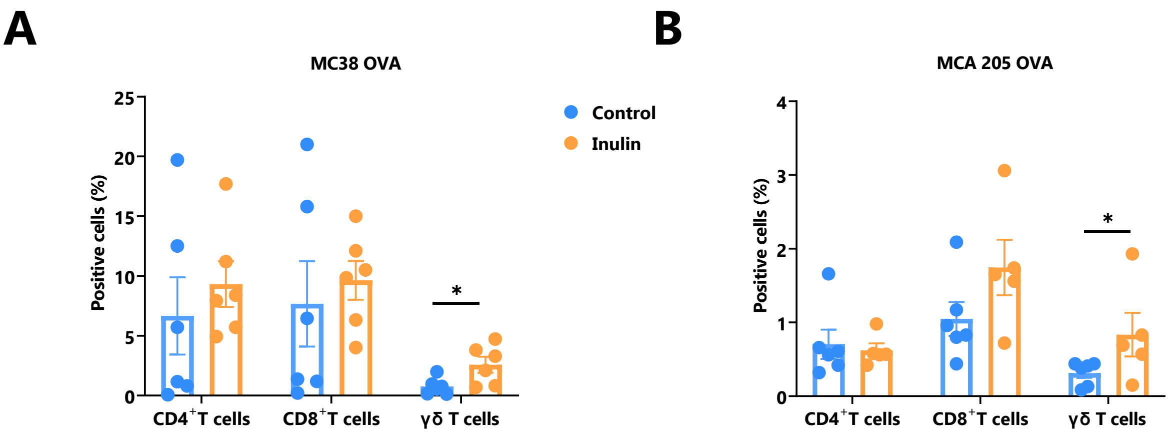


**Supplemental Figure S3: Inulin regimen promotes γδ T cell infiltration in MC38 OVA and MCA 205 OVA tumor models.**

C57BL/6 mice were fed with a control or an inulin-enriched diet (7.2% in drinking water) (n=6 mice per group) starting 15 days before subcutaneous (s.c.) inoculation of $5x10^5$ MC38 OVA colorectal cancer cells, or $2x10^5$ MCA 205 OVA fibrosarcoma cells (n=6 mice per group). Frequency of (A) MC38 OVA or (B) MCA 205 OVA tumor-infiltrated IFNγ-producing T lymphocytes. Graphs show the mean ± SEM. Statistically significant results are indicated by: *p < 0.05 by Mann-Whitney tests.

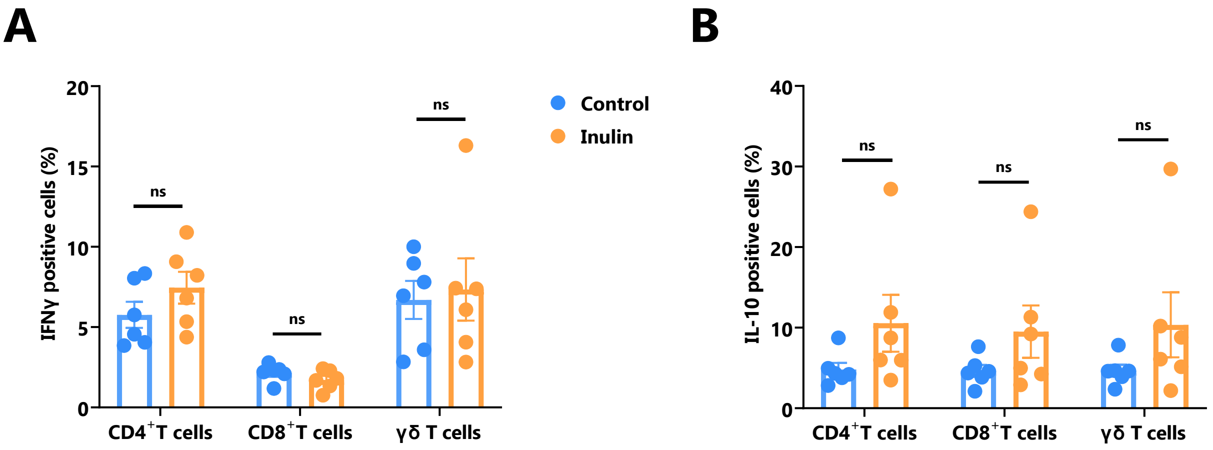


**Supplemental Figure S4: Inulin consumption tends to increase IL-10 production by LPC in the colon.**

C57BL/6 mice were fed with a control diet or inulin-enriched diet (7.2% in drinking water) (n=12 mice per group) for 15 days before the analysis of their gut immunity. Frequency of (A) IFNγ-producing or (B) IL-10-producing Lamina Propria T Cells (LPCs). Graphs show the mean ± SEM. ns = not-significant, *p < 0.05 by Mann-Whitney tests.

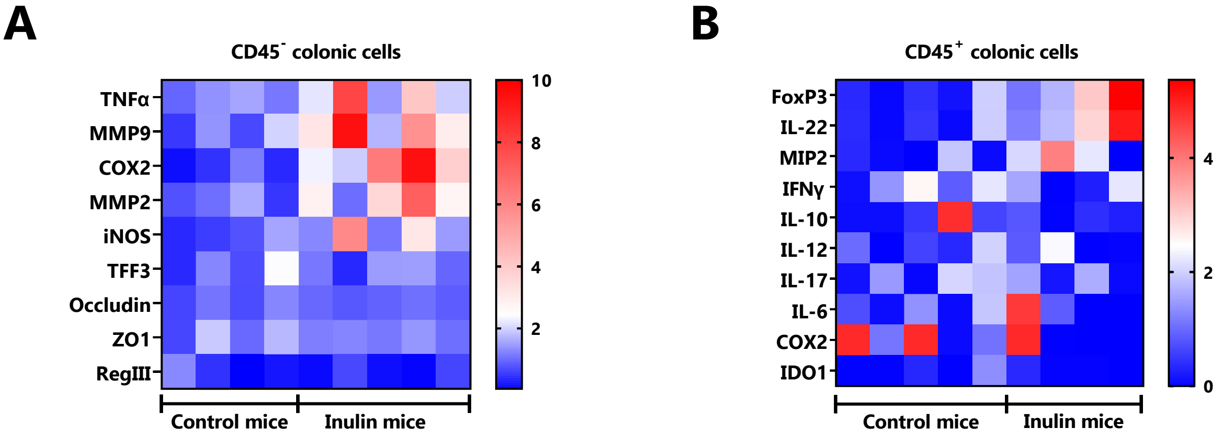


**Supplemental Figure S5: Inulin diet impacts immune and epithelial cell inflammation in the colon.**

qRT-PCR analysis of inflammation, tissue repair and tight junction -related genes in (A) $CD45^-$ and (B) $CD45^+$ colon cells sorted as described in (3E). Graphs show the expression levels of all the analysed genes.

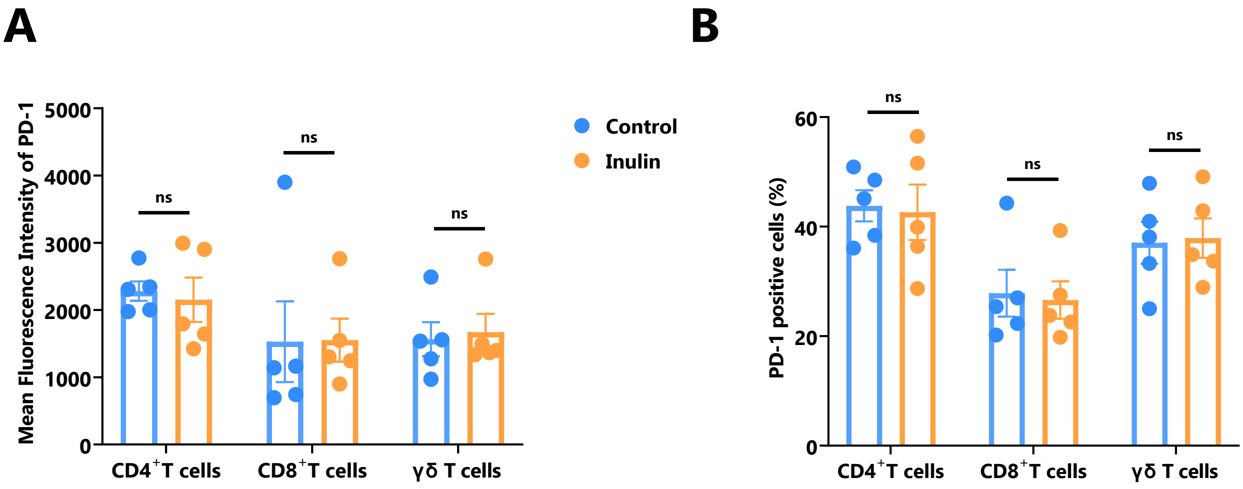


**Supplemental Figure S6: Inulin does not affect PD1 expression by TILs in the B16 OVA tumor.** (A) Mean Fluorescence Intensity (MFI) of PD1 and (B) frequency of PD1$^+$ cells among CD4$^+$, CD8$^+$ and γδ T cells in the B16 OVA tumor of mice treated as described in Figure 1A. Graphs show the mean ± SEM. ns = not-significant by Mann-Whitney tests.

| Percentages (%) | Control | Inulin | Global |
|---|---|---|---|
| **Firmicutes** | 59,61 | 57,11 | 58,30 |
| **Bacteroidota** | 37,87 | 35,87 | 36,83 |
| **Desulfobacterota** | 1,52 | 1,11 | 1,31 |
| **Proteobacteria** | 0,97 | 1,23 | 1,11 |
| **Actinobacteriota** | 0,03 | 4,67 | 2,46 |

**Supplemental Table S1: Inulin diet alters microbiota composition** Relative abundance of bacterial phyla in %. Means of 10-11 mice / group

| Antibody | Source | Identifier |
|---|---|---|
| APC/Cy7 anti-mouse CD45 | BD Bioscience | Cat# 557659, clone 30-F11, RRID:AB_396774 |
| BV421 anti-mouse PD-1 | BD Bioscience | Cat# 748268, Clone RMP1-30, RRID:AB_2872696 |
| BV421 anti-mouse IL-17A | BD Bioscience | Cat# 563354, Clone TC11-18H10, RRID:AB_2687547 |
| FITC anti-mouse CD3 | Biolegend | Cat# 100204, Clone 17A2, RRID:AB_312661 |
| PerCP/Cy5.5 anti-mouse CD4 | Biolegend | Cat# 100434, Clone GK1.5, RRID:AB_893324 |
| PE anti-mouse CCR9 | Biolegend | Cat# 129708, Clone 9B1, RRID:AB_2073249 |
| FITC anti-mouse CD11c | Biolegend | Cat# 117306, Clone N418, RRID:AB_313775 |
| PerCP/Cy5.5 anti-mouse CD8 | Biolegend | Cat# 100734, Clone 53-6.7, RRID:AB_2075238 |
| BV421 anti-mouse PD-L2 | Biolegend | Cat# 329616, Clone 24F.10C12, RRID:AB_2716087 |
| APC anti-mouse PD-L1 | Biolegend | Cat# 124312, Clone 10F.9G2, RRID:AB_10612741 |
| PerCP/Cy5.5 anti-mouse MHC Class II | Biolegend | Cat# 107626, Clone M5/114.15.2, RRID:AB_2191071 |
| PE/Cy7 anti-mouse CD80 | Biolegend | Cat# 104734, Clone 16-10A1, RRID:AB_2563113 |
| PE anti-mouse CD40 | Biolegend | Cat# 124610, Clone 3/23, RRID:AB_1134075 |
| APC anti-mouse IFNɣ | Biolegend | Cat# 505810, Clone XMG1.2 RRID:AB_315404 |
| PerCP/Cy5.5 anti-mouse CD11b | eBioscience | Cat# 45-0112-82, Clone M1/70, RRID:AB_953558 |
| PE/Cy7 anti-mouse ɣδ TcR | eBioscience | Cat# 25-5711-82, Clone eBioGL3 (GL-3, GL-3), RRID:AB_2573464 |
| PE anti-mouse IL-10 | Thermo Fisher Scientific | Cat# 12-7101-82, Clone JES5-16E3, RRID:AB_466176 |

**Supplemental Table S2: Antibodies used in flow cytometry.**

| Primer | Sequence (5' → 3') |
|---|---|
| GAPDH FW<br>GAPDH RV | GGTGAAGGTCGGTGTGAACG<br>CTCGCTCCTGGAAGATGGTG |
| β2M FW<br>β2M RV | GTATACTCACGCCACCCACC<br>TCCCGTTCTTCAGCATTTGG |
| IL-1β FW<br>IL-1β RV | TGATGAGAATGACCTCTTCT<br>CTTCTTCAAAGATGAAGGAAA |
| IL-6 FW<br>IL-6 RV | TAGTCCTTCCTACCCCAATTTCC<br>TTGGTCCTTAGCCACTCCTTCC |
| IL-10 FW<br>IL-10 RV | TCCTTAATGCAGGACTTTAAGGG<br>GGTCTTGGAGCTTATTAAAAT |
| IL-12 FW<br>IL-12 RV | CCTGGGTGAGCCGACAGAAGC<br>CCACTCCTGGAACCTAAGCAC |
| IL-17 FW<br>IL-17 RV | GCTCCAGAAGGCCCTCAGACTACC<br>CTTCCCTCCGCATTGACACAGC |
| TNF-α FW<br>TNF-α RV | AACTAGTGGTGCCAGCCGAT<br>CTTCACAGAGCAATGACTCC |
| IFNɣ FW<br>IFNɣ RV | GAACTGGCAAAAGGATGGTGA<br>TGTGGGTTGTTGACCTCAAAC |
| MIP-2 FW<br>MIP-2 RV | TCAATGCCTGAAGACCCTGC<br>CGTCACACTCAAGCTCTGGA |

| Primer | Sequence (5' → 3') |
|---|---|
| MMP-9 FW<br>MMP-9 RV | TGGGGGCAACTCGGC<br>GGAATGATCTAAGCCCAG |
| iNOS FW<br>iNOS RV | GTTGAAGACTGAGACTCTGG<br>ACTAGGCTACTCCGTGGA |
| Cox-2 FW<br>Cox-2 RV | GGGTTGCTGGGGGAAGAAATG<br>GGTGGCTGTTTTGGTAGGCTG |
| TFF-3 FW<br>TFF-3 RV | CCTGGTTGCTGGGTCCTCTG<br>GCCACGGTTGTTACACTGCTC |
| Occludin FW<br>Occludin RV | ACGGACCCTGACCACTATGA<br>TCAGCAGCAGCCATGTACTC |
| ZO-1 FW<br>ZO-1 RV | GGGGCCTACACTGATCAAGA<br>TGGAGATGAGGCTTCTGCTT |
| FoxP3 FW<br>FoxP3 RV | CCTATGGCTCCTTCCTTGGC<br>CCTTGGGTGCAGTCTTCCAG |
| RegIIIɣ FW<br>RegIIIɣ RV | TGGAGGTGGATGGGAATGGA<br>GCCACAGAAAGCACGGTCTA |
| IL-22 FW<br>IL-22 RV | GTGCTCAACTTCACCCTGGA<br>GGCTGGAACCTGTCTGACTG |
| IDO1 FW<br>IDO1 RV | TGGGACATTCCTTCAGTGGC<br>TCTCGAAGCTGCCCGTTCT |

**Supplemental Table S3: Sequence-specific primers used in qRT-PCR.** FW = Forward sequence; RV = Reverse sequence.
